# Structure of the triose-phosphate/phosphate translocator reveals the basis of substrate specificity

**DOI:** 10.1101/169169

**Authors:** Yongchan Lee, Tomohiro Nishizawa, Mizuki Takemoto, Kaoru Kumazaki, Keitaro Yamashita, Kunio Hirata, Ayumi Minoda, Satoru Nagatoishi, Kouhei Tsumoto, Ryuichiro Ishitani, Osamu Nureki

## Abstract

The triose-phosphate/phosphate translocator (TPT) catalyzes the strict 1:1 exchange of triose phosphate, 3-phosphoglycerate and inorganic phosphate across the chloroplast envelope, and plays crucial roles in photosynthesis. Despite rigorous studies for more than 40 years, the molecular mechanism of TPT is poorly understood due to the lack of structural information. Here we report crystal structures of TPT bound to two different substrates, 3-phosphoglycerate and inorganic phosphate, in occluded conformations. The structures reveal that TPT adopts a 10-transmembrane drug/metabolite transporter fold. Both substrates are bound within the same central pocket, where conserved lysine, arginine, and tyrosine residues recognize the shared phosphate group. A structural comparison with the outward-open conformation of the bacterial drug/metabolite transporter suggests a rocking-type motion of helix bundles, and molecular dynamics simulations support a model in which this helix rocking is tightly coupled to the substrate binding, to ensure strict 1:1 exchange. These results reveal the unique mechanism of sugar phosphate/phosphate exchange by TPT.

## Introduction

Photosynthetic organisms assimilate CO_2_ into organic compounds through the Calvin cycle^1^ and provide essential building blocks for all life on earth. In plants and algae, fixed carbons are exported from the chloroplast in the form of triose phosphate (triose-P)^2,3^. This export is accomplished by the triose-phosphate/phosphate translocator (TPT)^4-6^, which catalyzes the exchange of triose-P, 3-phosphoglycerate (3-PGA) and inorganic phosphate (Pi) across the chloroplast inner envelope membrane (Extended Data Fig. 1a).

**Figure 1.**
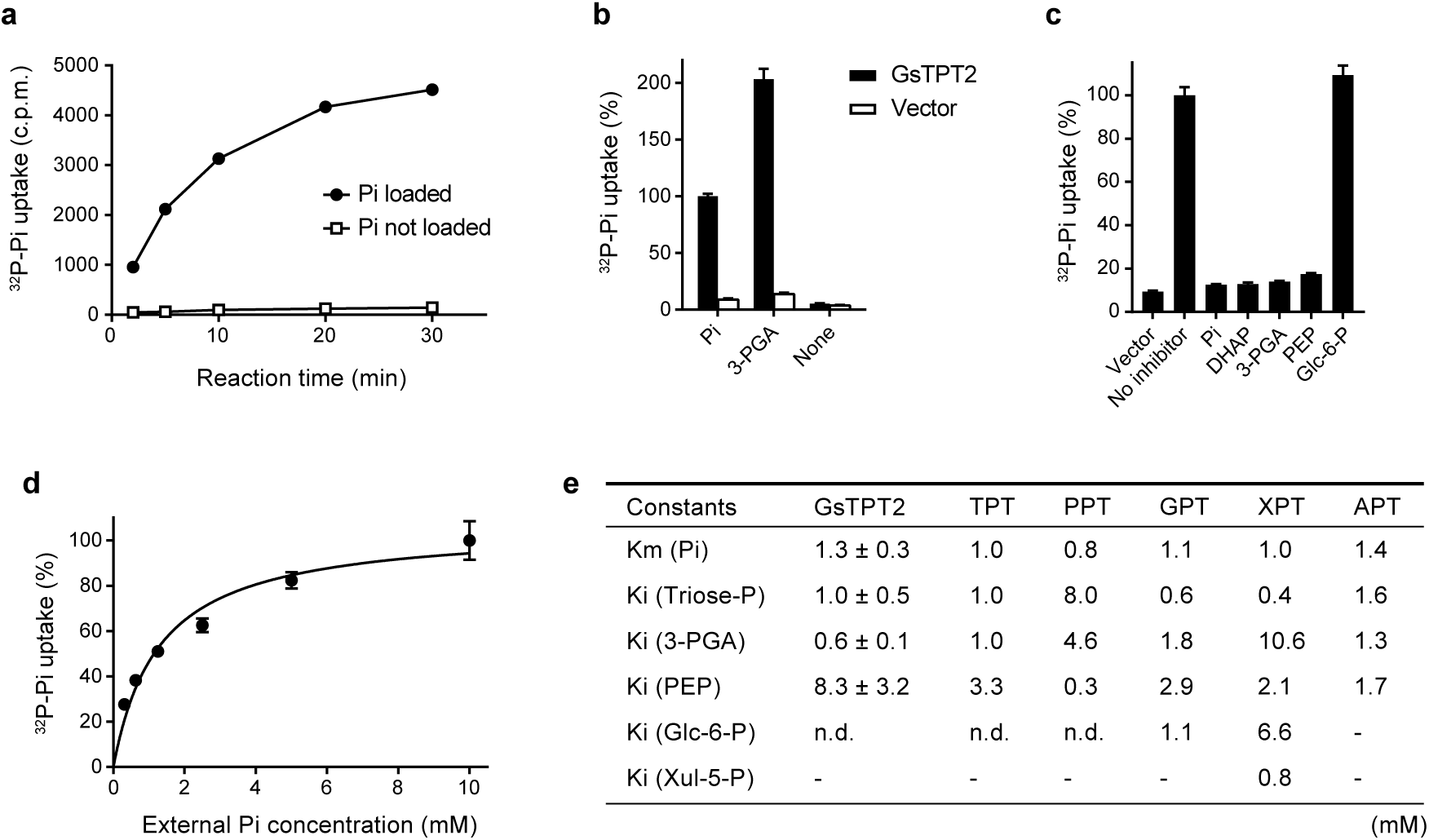
Functional characterization of GsTPT2. (a) Pi/Pi homo-exchange activity of GsTPT2. Liposomes were reconstituted with yeast membranes expressing GsTPT2, and the time-dependent uptake of [^32^P]-Pi was measured in the presence (filled circles) or absence (open squares) of internal Pi (30 mM). Error bars are s.e.m. (n=3). (b) Counter-flow assay. The uptake of [^32^P]-Pi was measured in the presence of the indicated internal substrates (30 mM). Control experiments were performed with membranes from yeast cells harboring empty vector. Error bars are s.e.m. (n=3). (c)Competitive inhibition assay. The uptake of [^32^P]-Pi was assayed in the presence of the indicated competitive inhibitors (40 mM) in the external solution. Error bars are s.e.m. (n=3). (d) Concentration-dependent uptake of [^32^P]-Pi. Error bars are s.e.m. (n=3). (e) Kinetic constants of GsTPT2 and the higher plant pPTs. The Michaelis constant (*K*_m_) of GsTPT2 was calculated from the experiment shown in Figure 1d. Inhibitor constants (*K*_i_) of GsTPT2 were evaluated at two different Pi concentrations with increasing inhibitor concentrations. Data are mean ± s.e.m. (n=3); n.d., not detectable. Values for the plant and apicomplexan pPTs were adopted from refs. ^9-11,18^.

TPT belongs to the plastidic phosphate translocator (pPT) family^7^, whose members are widespread across all photosynthetic eukaryotes^5^, as well as in other organisms with plastids^8^. Land plants possess four pPT subtypes, including the phosphoenolpyruvate/phosphate translocator (PPT)^9^, the glucose-6-phosphate/phosphate translocator (GPT)^10^ and the xylulose-5-phosphate/phosphate translocator (XPT)^11^, which transport different sugar phosphates and function in various metabolic pathways (Extended Data Fig. 1b, c). These plant pPTs play crucial roles in crop metabolism^12,13^, and therefore they are regarded as attractive targets of genetic manipulation for improving crop productivity^14,15^. The pPTs are also found in apicomplexan parasites^16,17^, which cause toxoplasmosis and malaria in humans. Since these apicomplexan pPTs are essential for the survival of the parasites^18,19^, they are potential drug targets for parasitic infections^20^. All pPT proteins catalyze strict 1:1 exchange reactions^3^, and thereby guarantee the total phosphate balance of the plastid and the cytosol while allowing the transport of carbon and energy^21^.

TPT was discovered more than forty years ago^3^ and has been rigorously characterized at genetic and biochemical levels^22^. However, due to the lack of structural information, the mechanisms by which TPT recognizes the substrates and catalyzes the strict 1:1 exchange remain poorly understood.

## Results

### Functional and structural analyses of TPT

To elucidate the structure and the molecular mechanism of TPT, we systematically screened plant and algal pPTs for their expression and stability. Among the tested proteins, a pPT from the thermophilic red alga *Galdieria sulphuraria* (Gasu_21660; Extended Data Fig. 2a,b) showed excellent solution behavior. Its crystallization construct (residues 91–410) without the N-terminal chloroplast transit peptide showed the ‘signature’ Pi/Pi homo-exchange activity^23^ in the liposome-based assay (Fig. 1a). We also confirmed the 3-PGA/Pi hetero-exchange activity (Fig. 1b). A competitive inhibition assay suggested that this pPT transports phosphorylated C3 compounds, but not phosphorylated C6 compounds (Fig. 1c). We further characterized its kinetic constants, and confirmed that its substrate specificity is comparable to that of higher plant TPTs: the Michaelis constant (*K*_m_) for Pi is about 1.3 mM, and the inhibition constants (*K*_i_) for triose-P, 3-PGA and phosphoenolpyruvate (PEP) are about 1.0, 0.6 and 8.3 mM, respectively (Fig. 1d,e). Based on these biochemical characterizations, we hereafter refer to this pPT as GsTPT2, although it was formerly named GsGPT^23^, based on the sequence similarity to GPT.

**Figure 2.**
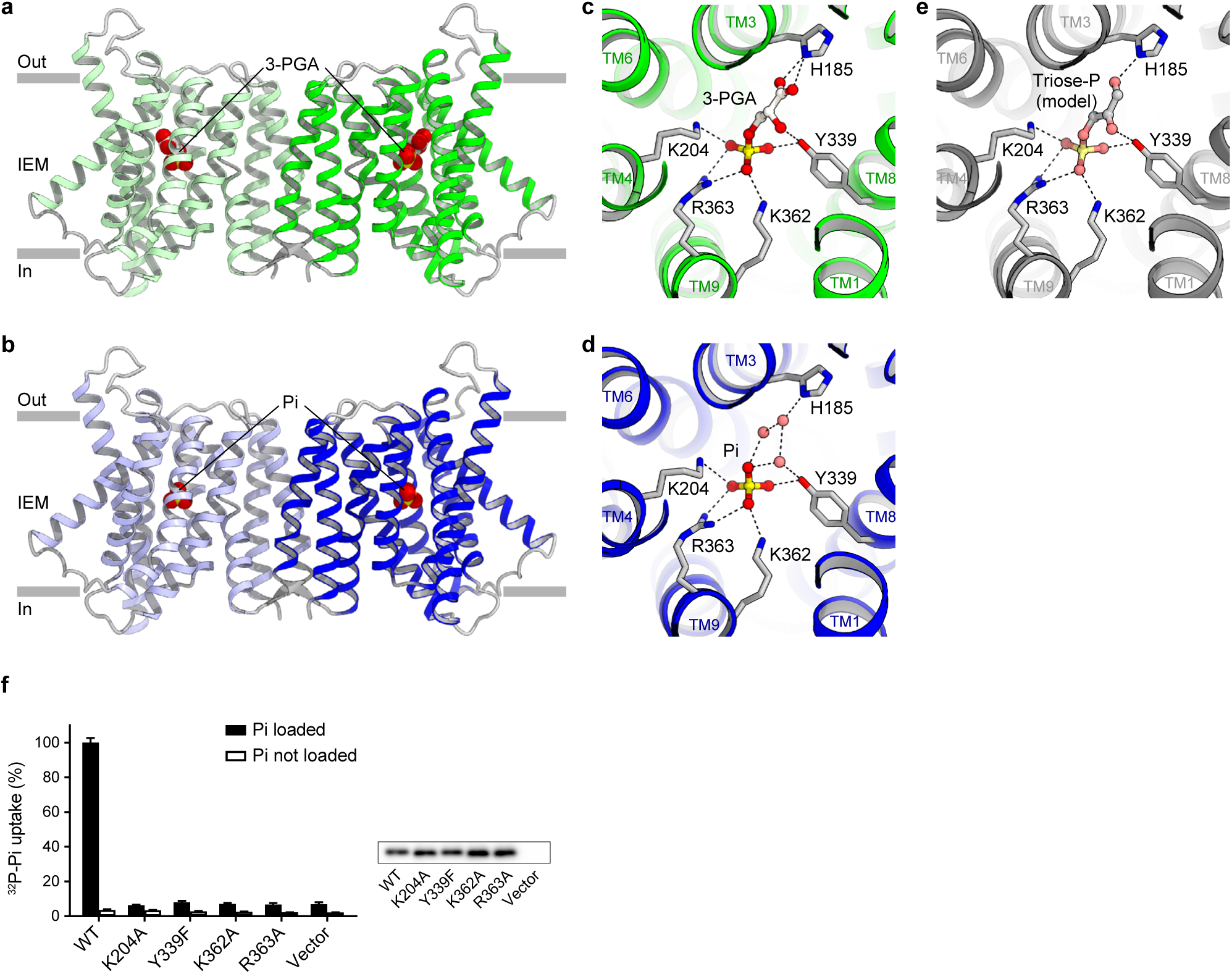
Overall structure and substrate recognition of GsTPT2. (a, b) Ribbon representations of the 3-PGA-bound (a) and the Pi-bound (b), occluded structures. IEM denotes the chloroplast inner envelope membrane. (c, d) Close-up view of the 3-PGA-binding site (c) and the Pi-binding site (d). Dotted lines indicate polar interactions. (e) Model of triose-P (dihydroxyacetone phosphate) binding. (f) Liposome-based mutational analysis. The levels of [^32^P]-Pi uptake by GsTPT2 mutants were compared to that of the wild-type. Error bars are s.e.m. (n=3). Western blotting confirmed the comparable expression levels of the wild type and mutant proteins (small inset).

We crystallized recombinant GsTPT2 in the lipidic cubic phase^24^. Co-crystallization with high concentrations (50–250 mM) of 3-PGA or Pi yielded diffraction-quality crystals belonging to the *P*2_1_2_1_2 space group, and we collected data from several hundred crystals, using a microfocus X-ray beam. A previous bioinformatics analysis suggested the classification of TPT into the drug/metabolite transporter (DMT) superfamily^25^, which contains various families of membrane proteins possessing 4, 5, 9 or 10 transmembrane (TM) helices. After extensive molecular replacement trials, we successfully obtained an initial solution with a poly-alanine model of the 10-TM DMT transporter SnYddG^26^. The final structures were determined at 2.1 and 2.2 Å resolutions for the Pi- and 3-PGA-bound states, respectively (Extended Data Figs. 3,4). The protein regions of the two structures are almost identical, with an r.m.s.d. value of 0.18 Å over 608 Cα atoms.

**Figure 3.**
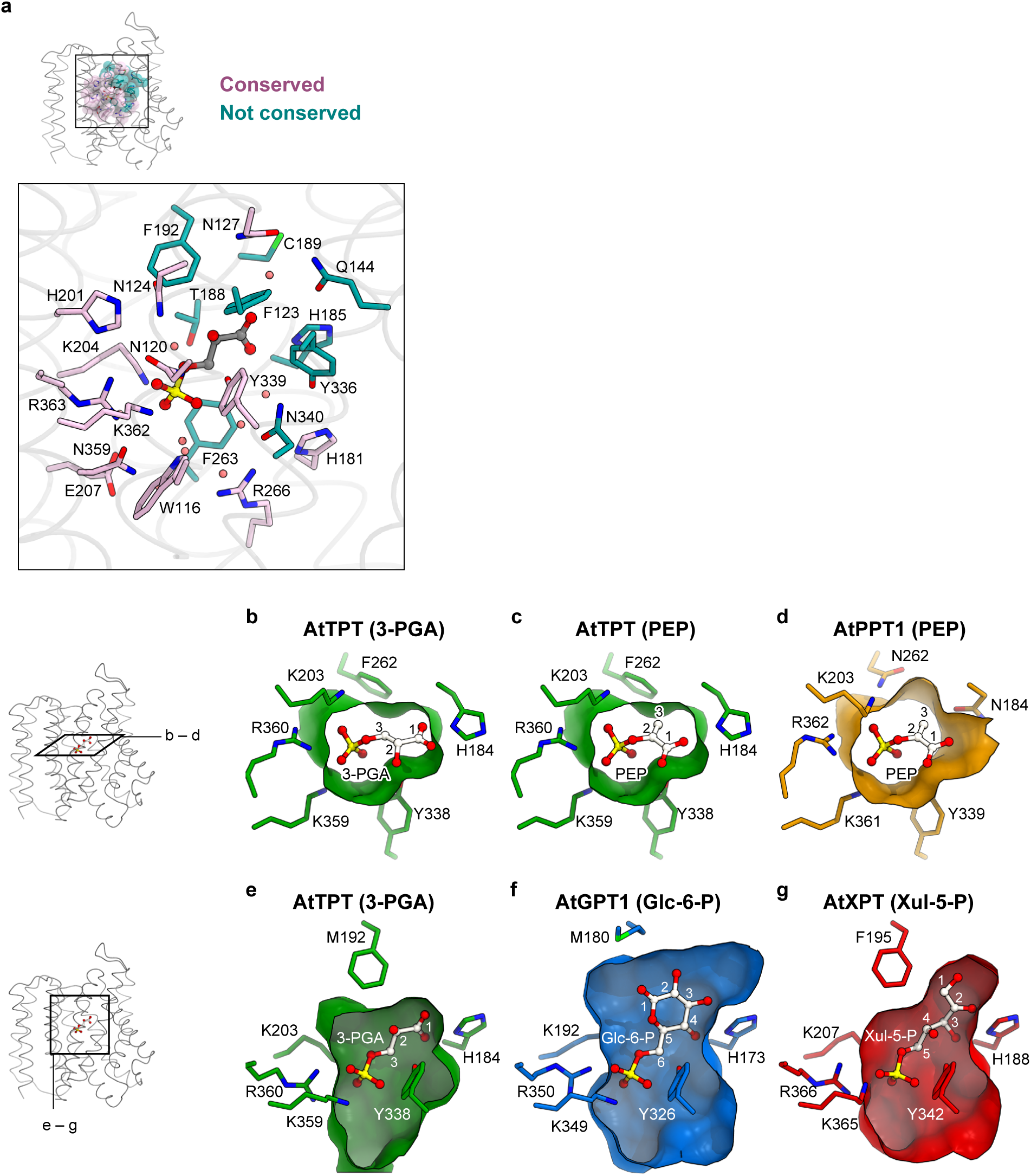
Comparison of the pPT subtypes. (a) Amino-acid sequence conservation of the substrate-binding site of GsTPT2. Conserved residues are colored violet and non-conserved residues are cyan. (b–g) Homology-modelled structures of the substrate-binding sites of the pPTs. Key residues involved in substrate recognition are shown as stick models. Substrate molecules were modelled manually, based on the coordination of 3-PGA in GsTPT2. Protein surfaces are shown for the regions around the substrate. In (c), the C3 carbon of PEP sterically clashed with the sidechain of Phe262 in the AtTPT model, indicating non-preferable binding.

**Figure 4.**
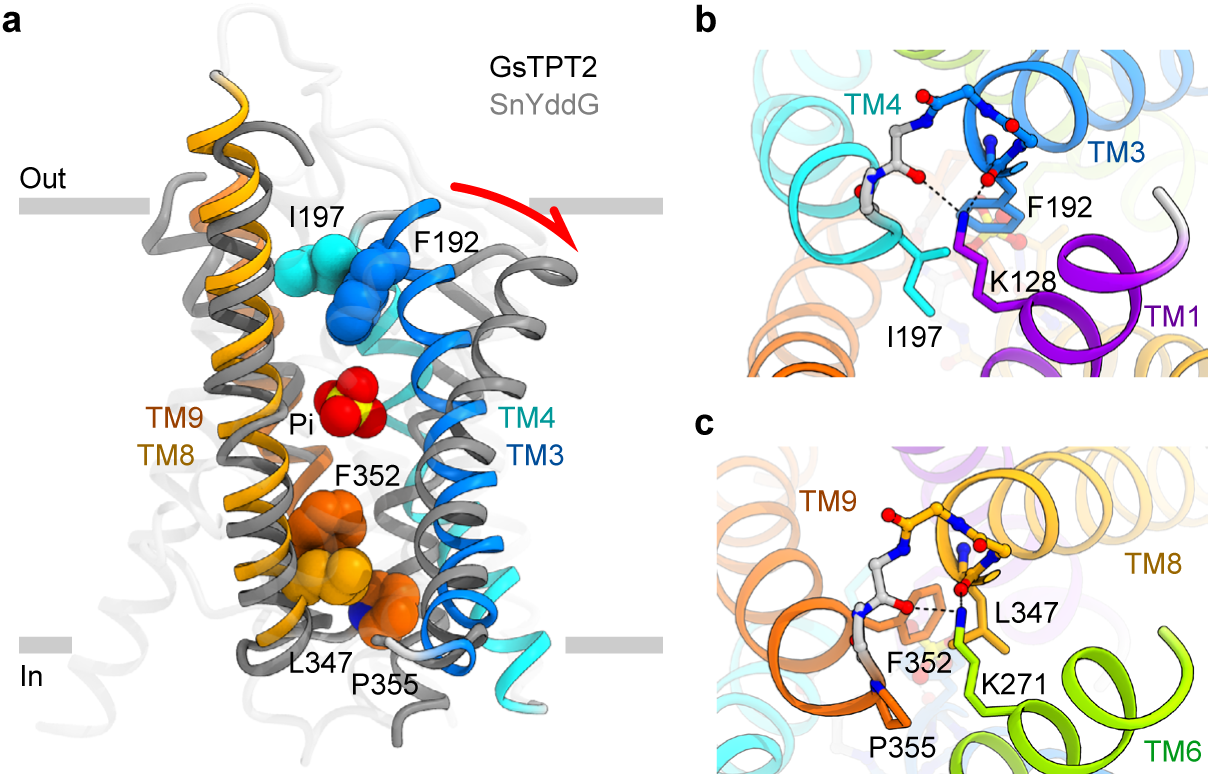
Occluded structure and alternating-access. (a) Superimposition of the occluded GsTPT2 structure and the outward-open SnYddG structure (PDB 5I20). The substrate and gate-forming residues are shown as CPK models. The red arrow highlights the putative rocking-type movements in TM3 and TM4. (b) Close-up view of the outside gate. (c) Close-up view of the inside gate.

The overall structure of GsTPT2 reveals a 10-TM helix topology with both the N- and C-termini on the stromal side (inside), rather than the previously predicted 6–9 TM helix topologies^27^ (Fig. 2a,b and Extended Data Fig. 5). GsTPT2 contains two ‘inverted’ structural repeats, comprising the N- and C-halves. Viewed from the intermembrane space side (outside), the five helices within the N- and C-halves (*i.e.*, TM1–5 and TM6–10) are arranged in counter-clockwise and clockwise manners, respectively (Extended Data Fig. 5). This fold is essentially similar to that of the bacterial DMT superfamily transporter SnYddG^26^, despite the low sequence identity (13.9%), suggesting that this “10-TM DMT fold” could be conserved across all putative 10-TM members of the DMT superfamily^25^ (Extended Data Fig. 6). In contrast to the ‘outward-open’ conformation of SnYddG, the current structure of GsTPT2 shows that its substrate-binding site is occluded from both sides of the membrane, revealing the ‘occluded’ conformation of a DMT protein for the first time.

**Figure 5.**
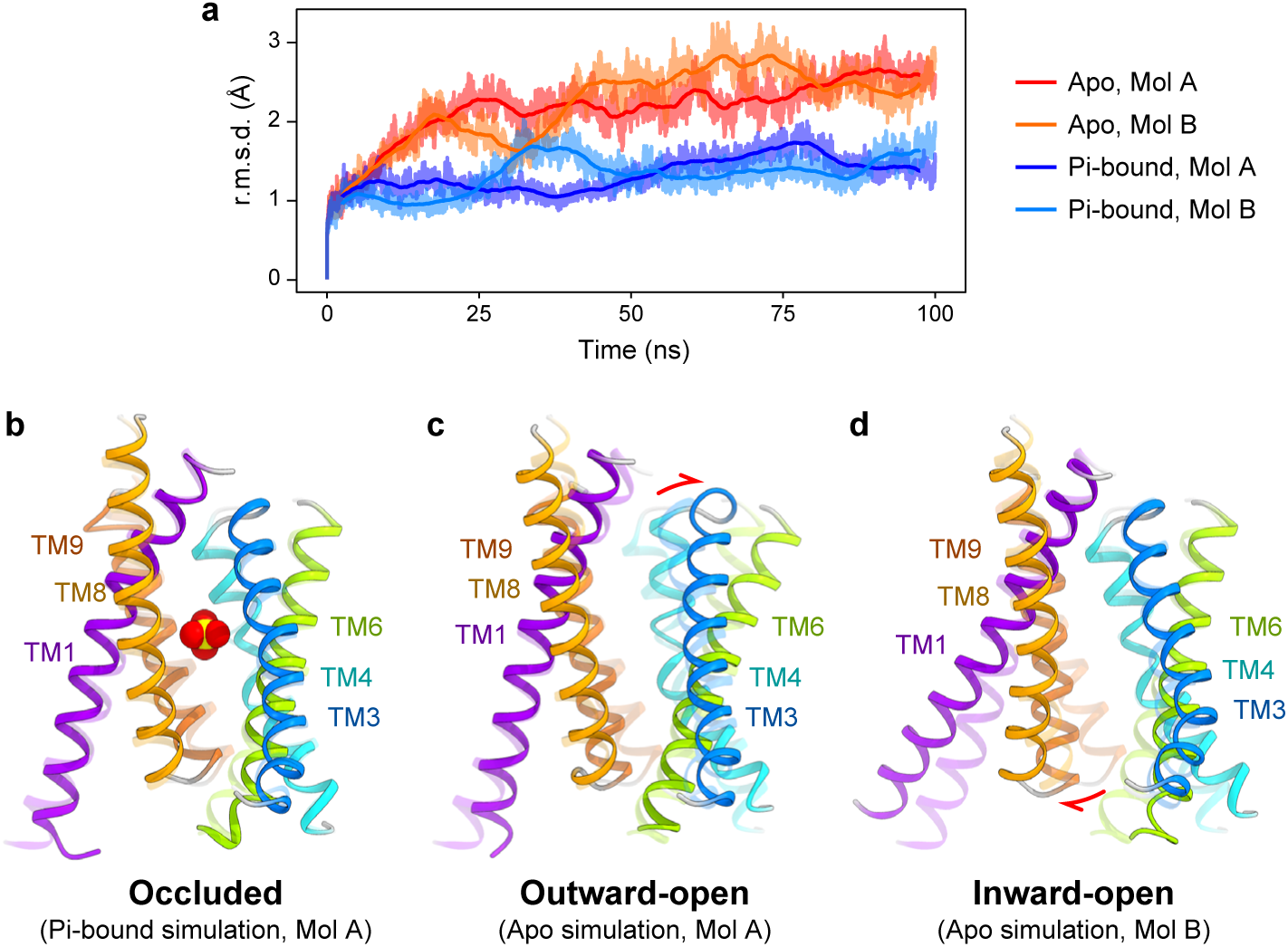
Molecular dynamics simulations. (a) R.m.s.d. plot of each monomer (Mol A and B) in the Pi-bound and apo simulations. (b–d) Comparison of the starting structures (0 ns, transparent) and the final structures (100 ns, opaque) in the simulations. Red arrows highlight helix movements.

**Figure 6.**
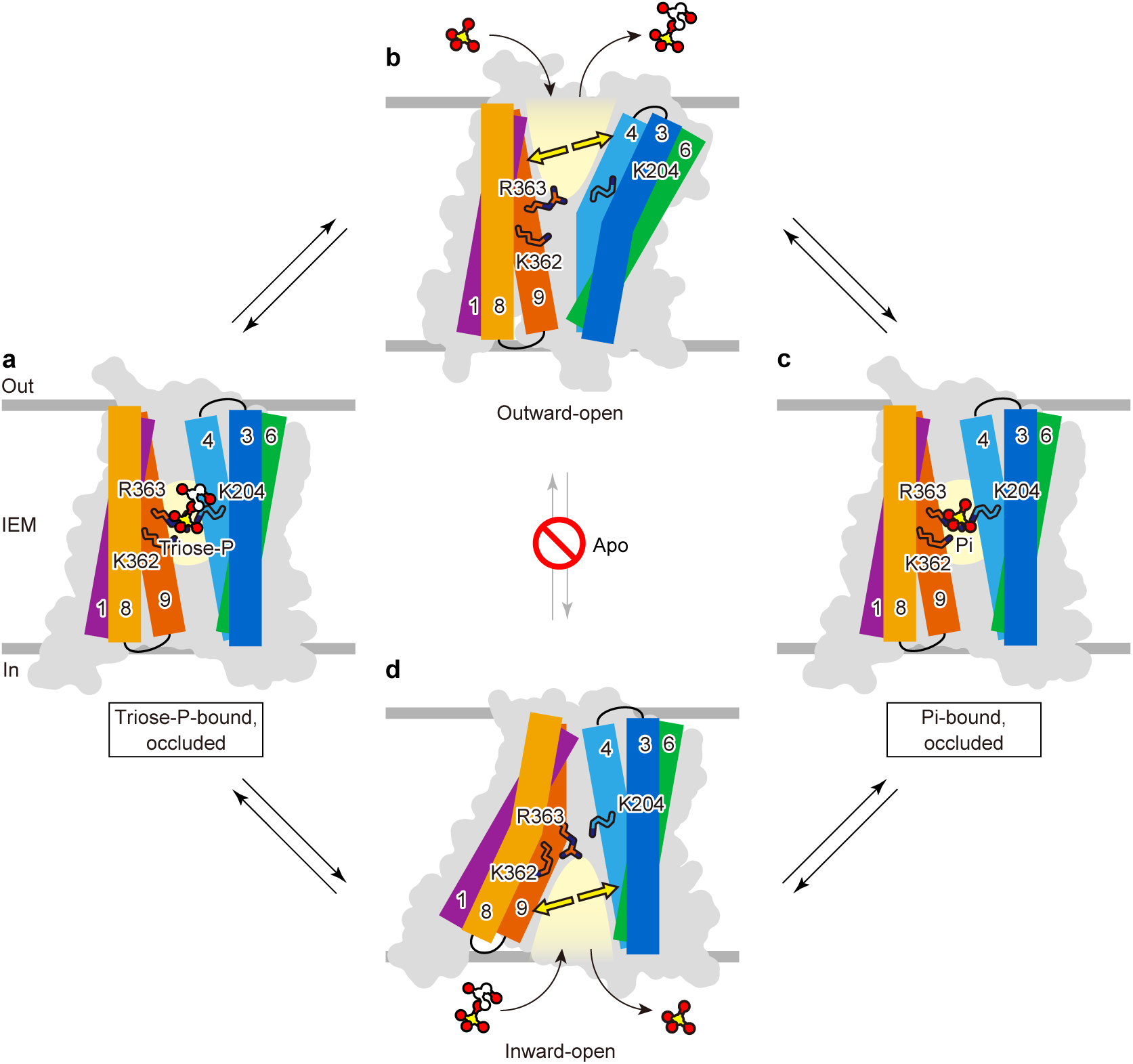
Proposed model for the strict 1:1 exchange. (a–d) Illustration of a hypothetical conformational cycle of GsTPT2. Substrate binding enables the association of the helix bundles (a, c), triggering the conformational transition. Without a substrate, the translocator cannot undergo the conformational transition due to the electrostatic repulsion (b, d).

Although the purified GsTPT2 protein is monomeric in solution, GsTPT2 forms a dimer in the crystallographic asymmetric unit (Extended Data Fig. 7). The inter-protomer interaction involves polar interactions at TM5, TM10 and a short β-strand connecting TM4 and TM5, and hydrophobic contacts through the lipid molecules bound at the interface. The same topological orientation of the monomers within the membrane suggests that this dimeric assembly could be physiologically relevant^28^, although we cannot exclude the possibility that this is a crystallization artifact.

### 3-PGA and Pi recognition

The electron density maps clearly showed that 3-PGA and Pi are bound to the same site located halfway across the membrane, as if trapped in a central ‘cage’ formed by TM1–4 and TM6–9 (Extended Data Fig. 8). The phosphate moiety of both ligands is identically recognized by ionic bonds with Lys204, Lys362 and Arg363 and a hydrogen bond with Tyr339 (Fig. 2c,d). While the three oxygen atoms of the phosphate (P-O2, O3 and O4) are directly recognized by these sidechains, the remaining oxygen atom (P-O1) is not directly recognized. In the 3-PGA-bound structure, this P-O1 is attached to the glycerate group, which extends into the space on the opposite side of the phosphate moiety and forms specific interactions with protein sidechains (Fig. 2c). The carboxyl group on the C1 atom forms an ionic bond with the sidechain of His185, and the hydroxyl group on the C2 atom hydrogen bonds with the sidechain of Tyr339. In addition, the C2 and C3 atoms form hydrophobic contacts with the sidechains of Thr188, Phe192 and Phe263 (Extended Data Fig. 8).

In the Pi-bound structure, the corresponding space near P-O1 is occupied by three water molecules (Fig. 2d). These water molecules form polar interactions with His185 and Tyr339, contributing to the indirect recognition of P-O1. Notably, the positions of these water molecules roughly correspond to those of the three oxygen atoms of the glycerate moiety of 3-PGA, mimicking the organic carbon structure. This water-mediated hydrogen-bonding network is likely to lower the energy of the Pi-bound state and could explain why Pi, which lacks a sugar moiety, is transported with a similar affinity to those of other sugar phosphates^21^.

The observed binding mode of 3-PGA suggests that triose-P can be recognized in a similar manner. Indeed, modelling of triose-P into the crystal structure indicates a good fit, with the oxygen atoms at the C1 and C2 positions forming similar polar interactions with His185 and Tyr339 (Fig. 2e). Therefore, the structures suggest that triose-P, 3-PGA and Pi, the three major counter-substrates of TPT, are similarly recognized in a single pocket. The structures also reveal that this pocket could not accommodate two or more phosphate moieties at a time, explaining why pyrophosphate or bisphosphate compounds are not readily transported across the chloroplast envelope membrane^3^.

### Similarity and diversity of pPT subtypes

The four phosphate-binding residues identified here are strictly conserved in all higher plant pPTs (Extended Data Fig. 2a). To examine their functional importance, we performed mutational assays of these residues (Fig. 2f). All of the tested mutations exhibited greatly reduced Pi/Pi homo-exchange activity, confirming their essential roles in phosphate transport. Besides these residues, a sequence comparison revealed that most of the residues near the phosphate moiety are also strictly conserved (Fig. 3a). In contrast, the residues distant from the phosphate, or near the sugar moiety, are varied among the different pPT subtypes (Fig. 3a). These findings suggest that the variant residues of the different pPT subtypes recognize the attached sugar moieties and thus determine their distinct substrate specificities, while the conserved residues similarly recognize the phosphate.

To better understand the substrate selectivities of the pPT family members^5^, we generated homology models of five representative pPTs, namely *Arabidopsis thaliana* TPT (AtTPT), PPT1 (AtPPT1), GPT1 (AtGPT1), and XPT (AtXPT) and *Toxoplasma gondii* APT (TgAPT). (Fig. 3 and Extended Data Fig. 9). The AtTPT model suggests that the plant TPTs similarly recognize the substrates as in the current GsTPT2 structure, since the residues recognizing 3-PGA are highly conserved (His185, Lys204, Tyr339, Lys362 and Arg363 in AtTPT) (Fig. 3b,e). TPT prefers three-carbon compounds phosphorylated at the C3 position (triose-P and 3-PGA) to those phosphorylated at the C2 position (PEP and 2-PGA) by ~10-fold^29^. The AtTPT model indicates that the ‘branched’ C3 methylene group of PEP would sterically clash with the bulky Phe262 sidechain, explaining the lower preference for PEP^9,30^ (Fig. 3c). In contrast, the C3 carbon of PEP can be accommodated in the widened pocket of the AtPPT1 model, where Phe is replaced by Asn262, consistent with the PPT’s preference for PEP^9^ (Fig. 3d). The apicomplexan pPTs, including TgAPT, PfipPT and PfopPT, have ‘dual specificity’, as they transport both triose-P and PEP with similar affinities^18,20^. The TgAPT model explains its dual specificity well, as it can accommodate both triose-P and PEP (Extended Data Fig. 7c,d).

GPT transports glucose-6-phosphate (Glc-6-P), the largest substrate of all pPTs, as well as smaller substrates such as triose-P and 3-PGA^10^. The AtGPT1 model has the largest pocket space, which can accommodate the bulky C6 sugar (Fig. 3f), consistent with its broad substrate specificity. As compared with GPT, the AtXPT model has a rather small pocket, which might be suitable for the C5 sugar moiety of its substrate, xylulose-5-phosphate^11^ (Xul-5-P) (Fig. 3g). Collectively, our crystal structures and the homology models address how different pPT members transport distinct sugar phosphates and thereby play diverse roles in plastid metabolism^5^.

### Basis of strict 1:1 exchange

Previous biochemical studies have shown that the transport by TPT is mediated by ‘alternating-access’^29^, in which the substrate-binding site is alternately exposed on either side of the membrane. In the current structure, the substrate is completely occluded from both sides of the membrane by the two gates (Fig. 4a). The ‘outside gate’ is formed by Phe192 and Ile197 on the tips of TM3 and TM4, and seals the substrate from the outside solvent (Fig. 4b). The ‘inside gate’ is formed by Leu347, Phe352 and Pro355 on the tips of TM8 and TM9, and similarly seals the substrate from the inside solvent (Fig. 4c). The helix ends of both gates are further capped by the conserved Lys128 and Lys271 residues (Fig. 4b,c).

To deduce the conformational change during the alternating-access, we compared this occluded structure with the available outward-open structure of the DMT transporter SnYddG. The structural superimposition revealed a prominent structural difference at TM3 and TM4 with a ~30° outward tilting in GsTPT2 (Fig. 4a), suggesting that these helices undergo rocking-type movements to open and close the outside gate. The pseudo-symmetric structure of GsTPT2 suggests that similar motion would occur in the symmetrical counterpart, TM8 and TM9, to open and close the inside gate (Supplementary Video 1).

To further understand the conformational changes, we performed molecular dynamics simulations of GsTPT2 in the presence or absence of the bound Pi (Fig. 5 and Extended Data Fig. 10). In the Pi-bound simulation, GsTPT2 did not undergo any significant structural change during 100 ns and remained in the occluded conformation (Fig. 5a,b). In contrast, in the apo simulation, GsTPT2 underwent rapid conformational changes within about 10 ns to the inward-open or outward-open conformations, and stably adopted these open conformations until the end of the simulation (~100 ns) (Fig. 5c,d and Supplementary Video 2). These conformational changes strongly support our model proposed from the structural comparison with SnYddG, which involves the rocking-type movement of the helix bundles TM3-TM4-TM6 and TM1-TM8-TM9 to open and close the two gates.

The different behaviors in the Pi-bound and apo simulations suggest that the conformational change of GsTPT2 is completely dependent on the substrate binding (Fig. 6). Without a substrate, due to the electrostatic repulsion between the cationic residues (Lys204, Lys362 and Arg363) in the middle of the helix bundles, GsTPT2 prefers the outward- or inward-open states, as shown in the MD simulation. In contrast, phosphate or organic phosphate binding allows the close approximation of these cationic residues and thus leads to the occluded state, as in the current crystal structures. This ligand-dependent conformational change ensures the substrate-dependent transition between the inward- and outward-open states, and thus explains the strict 1:1 exchange kinetics of the pPTs^21^.

The proposed coupling mechanism between the substrate binding and the conformational change is quite different from the transport mechanism proposed for YddG, another 10-TM member of the DMT superfamily. YddG is a uniporter^26^ that permeates substrates down a concentration gradient, indicating the lack of structural coupling. This difference could be explained by the composition of its substrate-binding site. The substrate-binding pocket of YddG mostly consists of hydrophobic residues^26^, which would lack electrostatic repulsion. YddG can thus adopt the occluded state without any substrate, consistent with its uniporter function. Therefore, even though GsTPT2 and YddG share the similar 10-TM DMT fold, the different compositions of their substrate-binding sites result in distinct transport mechanisms.

In conclusion, we determined the high-resolution structures of TPT in complex with two counter-substrates. The structures resolve the long-standing controversy over its helix topology^4,27^ and provide the framework to address its substrate recognition and strict 1:1 exchange mechanism. Further mechanistic understanding of the pPT family members could provide opportunities to engineer chloroplast transporters for improving crop productivity^14,15^, or to develop new drugs targeting plastid organelles of apicomplexan parasites^20^.

## Author contributions

Y.L. designed the research, expressed, purified and crystallized GsTPT2, determined the structures, and performed biochemical assays. Y.L., K.Y., and K.H. collected and processed diffraction data. T.N. and K.K. assisted with the structure determination. M.T. performed the molecular dynamics simulations. A.M. prepared *G. sulphuraria* cDNA. S.N. and K.T. performed the SEC-MALLS experiment. Y.L., T.N., R.I., and O.N. wrote the manuscript with help from all authors. O.N. directed and supervised all of the research.

## Acknowledgements

We thank H. Nishimasu for comments on the manuscript; A. Kurabayashi, K. Ogomori, W. Shihoya and R. Taniguchi for technical assistance; and the beam-line scientists at SPring-8 BL32XU for assistance in data collection. The diffraction experiments were performed at SPring-8 BL32XU (proposals 2015B0119 and 2015B2057). Computations of MD simulations were partially performed on the NIG supercomputer at ROIS National Institute of Genetics. This work was supported by grants from the Platform for Drug Discovery, Informatics and Structural Life Science by the Ministry of Education, Culture, Sports, Science and Technology (MEXT), JSPS KAKENHI (Grant Nos. 24227004, 25291011), the FIRST program, and a Grant-in-Aid for JSPS Fellows.

## Methods

### Cloning, expression and purification

Total cDNA from *G. sulphuraria* was prepared from autotrophically grown cells. The region of GsTPT2 (GI: 194462447) encoding residues 91–410 was amplified from the cDNA and subcloned into a modified pFastbac vector, with a C-terminal TEV cleavage site, EGFP and a His_10_-tag. Recombinant baculoviruses were produced with the Bac-to-Bac system (Invitrogen), and were used to infect Sf9 cells at a density of 2–3×10^6^ cells ml^−1^. After growth for 48 h at 27°C, the cells were harvested and sonicated in lysis buffer (50 mM Tris-HCl (pH 8.0), 150 mM NaCl and protease inhibitors). The cell debris was removed by low-speed centrifugation (10,000 g, 10 min), and the membrane fraction was collected by ultracentrifugation (138,000 g, 1 h).

The membrane fraction was solubilized in solubilization buffer (20 mM Tris-HCl (pH 8.0), 300 mM NaCl, 1% (w/v) lauryl maltoside neopentyl glycol (LMNG) and 1 mM β-mercaptoethanol (β-ME)) for 3 h at 4°C. The supernatant was isolated by ultracentrifugation (138,000 g, 30 min) and subjected to immobilized metal ion affinity chromatography (IMAC) with Ni-NTA resin (Qiagen). The resin was washed with IMAC buffer (20 mM Tris-HCl (pH 8.0), 300 mM NaCl, 0.05% LMNG, 1 mM β-ME and 30 mM imidazole), and the protein was eluted with IMAC buffer supplemented with 300 mM imidazole. The eluate was treated with TEV protease and dialyzed overnight against dialysis buffer (20 mM Tris-HCl (pH 8.0), 300 mM NaCl, 0.01% LMNG and 1 mM β-ME). The cleaved EGFP-His_10_ and TEV protease were removed by reverse IMAC with Ni-NTA. The protein was concentrated to 2–3 mg ml^−1^ using a 50 kDa MWCO concentrator (Millipore), and further purified by size-exclusion chromatography (SEC) on a Superdex 200 Increase 10/300 column (GE Healthcare) in SEC buffer (10 mM Tris-HCl (pH 8.0), 150 mM NaCl, 0.01% LMNG and 1 mM β-ME). The peak fractions were collected, concentrated to 10–20 mg ml^−1^, flash-frozen in liquid nitrogen and stored at −80°C until crystallization.

### Size exclusion chromatography coupled to multi-angle laser light scattering (SEC-MALLS)

The instrument setup for the SEC-MALLS experiment consisted of an Agilent 1100 Series HPLC system connected in series with a Shimadzu SPD-10Avp UV absorbance detector, a Wyatt DAWN HELEOS 8+ light scattering detector and a Shodex RI 101 refractive index detector. Analytical size-exclusion chromatography was performed at 25 °C on a Superdex 200 10/300 column equilibrated with buffer containing 10 mM Tris-HCl (pH 8.0), 150 mM NaCl and 0.01% LMNG. A 90 μl portion of the purified GsTPT2 sample (1.5 mg ml^−1^) was injected into the column and eluted at 0.5 ml min^−1^. Elution was monitored in line with the three detectors, which simultaneously measured UV absorption, light scattering and refractive index. A 658 nm laser was used in the light scattering measurement. Molecular masses were calculated using the three-detector method^31^, as implemented in the ASTRA software package (Wyatt Technology).

### Crystallization

Purified samples were thawed and mixed with 1-oleoyl-R-glycerol (monoolein), at a protein to lipid ratio of 2:3 (w/w), to prepare the lipidic cubic phase (LCP), as previously described^32^. Crystallization experiments were performed with 96-well glass sandwich plates (Molecular Dimensions), using a Gryphon LCP robot (Art Robbins Instruments). Typically, 50 nl of protein-laden LCP drops were overlaid with 800 nl of precipitant solution. After extensive co-crystallization screening, needle-shaped crystals appeared under conditions containing high concentrations of 3-PGA or Pi. Optimized crystals of the 3-PGA-bound state were obtained in 35–40% PEG200, 100 mM Na-citrate (pH 6.0), 50–100 mM citrate 3K and 50–100 mM 3-PGA·2Na. Optimized crystals of the Pi-bound state were obtained in 43–48% PEG200, 50–100 mM MES-NaOH (pH 6.0) and 200–250 mM (NH_4_)_2_HPO_4_. Crystals were harvested and flash-cooled in liquid nitrogen for data collection.

### Data collection and structure determination

X-ray diffraction experiments were performed at the micro-focus beamline BL32XU at SPring-8. The locations of well-diffracting crystals were identified by raster scanning, and data were collected for a 5–30° wedge from each crystal. All diffraction data were processed with XDS^33^, and merged with XSCALE based on the hierarchical clustering analysis with BLEND^34^ or with the cross-correlation method as implemented in the KAMO software (https://github.com/keitaroyam/yamtbx).

For the determination of the Pi-bound structure, we performed molecular replacement trials with Phaser^35^ using various truncated structures of DMT proteins as search models. Initial solutions were obtained with a full-length polyalanine model of the YddG monomer (PDB 5I20). After initial refinement in PHENIX^36^, the resulting map (Rfree value 53%) showed poor or no electron density for substantial portions of the structure, particularly for TM5, TM10 and all loop regions. These invisible segments were deleted from the model and rebuilt by multiple trials of manual modelling of new polyalanine helices using COOT^37^ and refinement with phenix.refine, to find the correct helix assignment. After subsequent rounds of model building and refinement, we could build ten helix backbones and several sidechains into the visible electron density. However, at this point, further model building did not improve the *R*_free_ value or the quality of the electron density. We then noticed that the model was of a ‘swapped’ form of the protein, where the N-terminal repeat (TM1–5) and the C-terminal repeat (TM6–10) were inversely assigned to each other. We corrected this swapping by renumbering the residues in COOT and proceeded with further model building. After building the protein regions (residues 100–404), strong electron densities were observed within the central cavities of the two monomers, which were unambiguously assigned as bound Pi molecules. During the later stages of refinement, electron densities for water and lipid molecules were also identified. The structure was iteratively rebuilt and refined with COOT and PHENIX to achieve good stereochemistry and *R*_free_ values (Extended Data Table 1). The 3-PGA-bound structure was determined using the Pi-bound GsTPT2 dimer as the starting model, and iteratively rebuilt and refined with COOT and PHENIX.

### Preparation of reconstituted liposomes

Yeast membranes expressing recombinant proteins were prepared as previously described^23^, with slight modifications. The region of GsTPT2 encoding residues 91–410 was cloned into a modified pYES2 vector, with a C-terminal His_6_ tag. For mutant assays, mutations were introduced by a PCR-based method. The plasmids were transformed into *Saccharomyces cerevisiae* cells (strain BY4742). Transformed cells were grown in CSM-URA medium containing 2% raffinose, and protein expression was induced with 2% galactose when the culture reached an OD_600_ = 0.6. After growth for 22 h at 30°C, the cells were harvested and disrupted in lysis buffer (50 mM Tricine-KOH (pH 7.5), 0.1 mM phenylmethylsulfonyl fluoride and 5% glycerol), using acid-washed glass beads (200–400 μm; Sigma). Glass beads and cell debris were removed by low-speed centrifugation (4,000 g, 2 min), and the membrane fraction was collected by ultracentrifugation (138,000 g, 1 h). The membrane pellet was resuspended in 50 mM Tricine-KOH (pH 7.5), flash-frozen in liquid nitrogen and stored at −80°C until use. Aliquots of the resuspended membranes were subjected to SDS-PAGE, and the His_6_-tagged recombinant proteins were detected by a western-blot analysis, using an anti-His-tag polyclonal antibody (code PM032; MBL).

Soybean L-α-phosphatidylcholine (Avanti) in chloroform was dried into a thin film under a stream of nitrogen gas, and further dried under vacuum. Dried lipids were resuspended at 20 mg ml^−1^ in intra-liposomal solution (120 mM Tricine-KOH (pH 7.5) and 30 mM NaH_2_PO_4_) or Pi-free intra-liposomal solution (150 mM Tricine-KOH (pH 7.5)), and sonicated for 5 min at 4°C to form unilamellar vesicles. This unilamellar vesicle solution was reconstituted with the yeast membranes at 19:1 (v/v), by the freeze-thaw procedure. The reconstituted liposomes were sonicated again for 5 min at 4°C, to form unilamellar vesicles. The extra-liposomal solution was exchanged by gel-filtration on Sephadex G-50 (GE Healthcare) pre-equilibrated with 150 mM Tricine-KOH (pH 7.5).

### Liposome assays

The liposome assays were performed as previously described^23^, with slight modifications.

For the time-dependent uptake assay, the reaction was started by mixing the reconstituted liposome solution with an equal volume of extra-liposomal solution (150 mM Tricine-KOH (pH 7.5) and 1 mM [^32^P]-NaH_2_PO_4_ (0.1 mCi ml^−1^)). At different time points, liposomes were isolated by anion exchange on AG-1 X8 resin (acetate form, 200–400 dry mesh size; Bio-Rad), pre-equilibrated with 150 mM sodium acetate. The radioactivity of the incorporated [^32^P]-Pi was quantified by liquid scintillation counting. Mutant assays were performed with a similar procedure, and the total amounts of incorporated [^32^P]-Pi were compared at 30 min.

For the counter-flow assay, the liposomes containing 30 mM Pi, 30 mM 3-PGA or no substrate were mixed with extra-liposomal solution containing 0.25 mM [^32^P]-Pi. The total amounts of incorporated [^32^P]-Pi were compared at 3 min.

For the competitive inhibition assay, the liposomes containing 30 mM Pi were mixed with extra-liposomal solution containing 0.25 mM [^32^P]-Pi and 40 mM of the indicated competitive inhibitor. The total amounts of incorporated [^32^P]-Pi were compared at 3 min.

For the determination of kinetic constants, the Michaelis constant (*K*_m_) for Pi was analyzed using various external concentrations of [^32^P]-Pi (0.315–10 mM) and a fixed internal concentration of Pi (30 mM). Inhibitor constants (*K*_i_) were assessed with two different external concentrations of [^32^P]-Pi (0.5–2.5 mM) and four different concentrations of the indicated inhibitors (0–10 mM). To assess the background uptake, control experiments were performed with membranes from yeast cells transformed with empty vector. Enzyme kinetic data were analyzed by non-linear regression fitting, as implemented in the GraphPad Prism 7 software.

### Molecular dynamics simulation

The simulation system included the GsTPT2 dimer, 1-phosphoryl-2-oleoylphosphatidylcholine (POPC), TIP3P water and 150 mM NaCl. The disordered sidechains in the GsTPT2 crystal structure were modelled by COOT^37^. To embed the protein within the POPC bilayer, the protocol described by Javanainen^38^ was used. One POPC molecule was placed in the GsGPT dimerization interface, corresponding to the two monoolein molecules in the crystal structure. Finally, the periodic boundary systems, including 136,668 (with Pi) and 136,652 (without Pi) atoms, with the size of 90.7×147.9×100.0 Å, were prepared. The net charge of the solute was neutralized with sodium and chloride ions. The molecular topologies and force field parameters from CHARMM36 (ref. ^39^) were used. Molecular dynamics simulations were performed by the program Gromacs, version 5.0.5 (ref. ^40^). First, energy minimization was performed using the steepest descent, with a cut-off of 1,000.0 kJ mol^−1^ nm^−1^. Next, random velocities were assigned according to a Maxwell distribution, at a temperature of 310 K for each atom, and an equilibration run (eq1) was performed for 100 ps in the canonical (*NVT*) ensemble (310 K, 90.7×147.9×100.0 Å volume). Finally, an equilibration run (eq2) was performed for 1,000 ps in the isothermal-isobaric (*NPT*) ensemble (310 K, 1 bar). The positions of non-hydrogen atoms in the protein and phosphates were restrained with a force constant of 1,000 kJ mol^−1^ nm^−2^, in the minimization and equilibration runs. Production runs were performed for 100 ns in the NPT ensemble (310 K, 1 bar). The same simulation was performed twice with different initial velocities, and similar results were obtained. Constant temperature was maintained by using V-rescaling^41^ with a time constant of 0.1 ps in eq1, and a Nosé-Hoover thermostat^42,43^ with a time constant of 0.5 ps in eq2 and the production runs. Pressure was controlled with semiisotropic coupling to a Parrinello-Rahman barostat^44^, with a time constant of 5.0 ps and a compressibility of 4.5×10^−5^ bar^−1^. The LINCS algorithm^45^ was used for bond constraints. Long range electrostatic interactions were calculated with the particle mesh Ewald method^46^.

## Extended Data

**Extended Data Figure 1 | Roles of the pPTs in plastid metabolism**

(a) The function of TPT in chloroplasts. Under physiological conditions, TPT catalyzes either triose-P/Pi or triose-P/3-PGA exchange across the chloroplast inner envelope membrane. The former reaction delivers a carbon skeleton to the cytoplasm and transports Pi back into the chloroplast for ATP regeneration. The latter reaction, known as the triose-P/3-PGA shuttle^2^, indirectly exports chemical energy (ATP and NADPH) without the net transport of carbon. OEM and IEM denote the outer and inner envelope membranes, respectively.

(b) The functions of other pPT subtypes. PPT exchanges PEP with Pi and plays important roles in amino acid and fatty acid syntheses^9^. PPT is also a part of the CO_2_ concentration mechanism of C4 and CAM photosynthesis^47^. GPT and XPT exchange Glc-6-P and Xul-5-P with Pi, respectively, and they play important roles in the starch biosynthesis and pentose-phosphate pathways^10,11^.

(c) Chemical structures of the pPT substrates.

**Extended Data Figure 2 | Sequence alignment of the pPTs**

(a) Amino acid sequence alignment of the pPTs.

(b) Sequence identity matrix of the pPTs. Identities were calculated for the mature

(a) translocator regions (corresponding to residues 101–410 in GsTPT2).

**Extended Data Figure 3 | Crystallization, data collection and structure determination of GsTPT2**

(a) Co-crystals of GsTPT2 and 3-PGA.

(b) Diffraction image of the 3-PGA-bound state.

(c) Magnified image of the region enclosed by the square in (b).

(d) Co-crystals of GsTPT2 and Pi.

(e) Diffraction image of the Pi-bound state.

(f) Magnified image of the region enclosed by the square in (e).

(g) Crystal packing of the 3-PGA-bound structure.

(h) Crystal packing of the Pi-bound structure.

*(i) F*o-*F*c omit map of 3-PGA, contoured at 4.0 σ.

*(j) F*o-*F*c omit map of Pi, contoured at 4.0 σ.

(k) Superimposition of the monomers in the 3-PGA-bound structure.

(l) Superimposition of the monomers in the Pi-bound structure

(m) Superimposition of the Pi-bound and 3-PGA-bound dimers.

(a) **Extended Data Figure 4 | Electron density map**

(a) 2*F*o-*F*c map (contoured at 1.7 σ) of the 3-PGA-bound structure. TM1, 8, and 9 are not shown for clarity.

(b) 2*F*o-*F*c map (contoured at 1.7 σ) of the Pi-bound structure.

**Extended Data Figure 5 | Structural pseudo-symmetry of GsTPT2**

(a) Transmembrane topology of GsTPT2. TM1 to TM10 are color-coded in rainbow. The asterisk highlights the position of the substrate-binding site.

(b) Superimposition of the two structural repeats. The N-terminal repeat (residues 100–245) and the C-terminal repeat (residues 246–404) were superimposed by the SSM algorithm^48^. TM5 and TM10 did not superimpose well, indicating local asymmetry.

**Extended Data Figure 6 | Structural similarity of the two DMT superfamily transporters**

(a) Sequence comparison between GsTPT2 and SnYddG. The two sequences were aligned by using Clustal omega^49^ and manually adjusted based on the secondary structure.

(b, c) Stereo views of the superimposition between GsTPT2 and SnYddG. The outer halves of TM3, TM4 and TM6 show structural differences relevant to the proposed gating mechanism. TM5 and TM10 also show large shifts, which are not relevant to the gating mechanism. The structures were superimposed by the SSM algorithm^48^. TM1–4 and TM6–9 were used for secondary structure element calculations, and 168 Cα atoms were used for the final three-dimensional alignment.

**Extended Data Figure 7 | Dimerization of GsTPT2 in the crystal**

(a) 2*F*o-*F*c map (contoured at 1.0 σ) of the six monoolein molecules identified near the dimer interface.

(b) Lipid-protein interactions. Four monoolein molecules mediate the hydrophobic contacts between the two protomers.

(c, d) Protein-protein interactions. Polar sidechains form hydrogen bonds between the protomers (c). The loops connecting TM4 and TM5 form a short, two-stranded β-sheet between the two protomers (d). Dotted lines indicate polar interactions.

(e) SEC-MALLS analysis of GsTPT2. The three chromatograms show the readings of the UV absorption, refractive index and light scattering detectors. The traces were normalized to the peak maxima. The cyan and red curves in the light scattering chromatogram indicate the calculated molecular masses of the protein-detergent complex and the protein, respectively (Mc and Mp; values are given on the right axis). The black arrow highlights the position of the elution peak of GsTPT2. The refractive index increments (dn/dc) of the protein and the detergent were assumed to be 0.185 and 0.132, respectively^50^.

(f) Molecular mass values determined by the SEC-MALLS experiment or calculated from the amino acid sequence. The experimental protein mass was determined to be about 29.6 kDa, corresponding to the theoretical mass of the GsTPT2 monomer, 36.4 kDa.

**Extended Data Figure 8 | Substrate recognition**

(a) Stereo view of the 3-PGA binding site. Dotted lines indicate polar interactions. Red balls represent water molecules.

(b) Stereo view of the Pi binding site.

(c) Schematic diagram of the 3-PGA coordination.

**Extended Data Figure 9 | Structural basis of substrate specificity in the pPT subtypes**

(a–g) Homology-modelled structures of the substrate-binding sites of the pPTs. In (b-g), the substrate molecules were modelled manually, based on the coordination of 3-PGA in GsTPT2 (a). Key residues involved in substrate recognition are shown as stick models. Protein surfaces are shown for selected regions around the substrates. In (b), the C3 carbon of the PEP model sterically clashed with the Phe263 sidechain in GsTPT2.

**Extended Data Figure 10 | Molecular dynamics simulations**

(a–d) Cut-away surface representations of the final structures (100 ns) in the simulations. We performed the apo and Pi-bound simulations independently twice. In the Pi-bound simulations, the translocator stably adopted the occluded conformation (a,b). In contrast, in the apo simulations, the two protomers adopted the inward- and outward-open conformations (c), or both adopted the inward-open conformation (d), suggesting that the presence or absence of the substrate in the central pocket significantly affected the translocator conformation, and supporting the substrate-dependent conformational change (discussed in the main text). In addition, the weak structural coupling of the two protomers (‘inward-outward’ or ‘inward-inward’) suggested that each protomer acts as an independent functional unit, consistent with the ping-pong type kinetics^51^. (e–l), Comparison of the starting structures (0 ns, transparent) and the final structures (100 ns, opaque).

**Extended Data Table 1.**
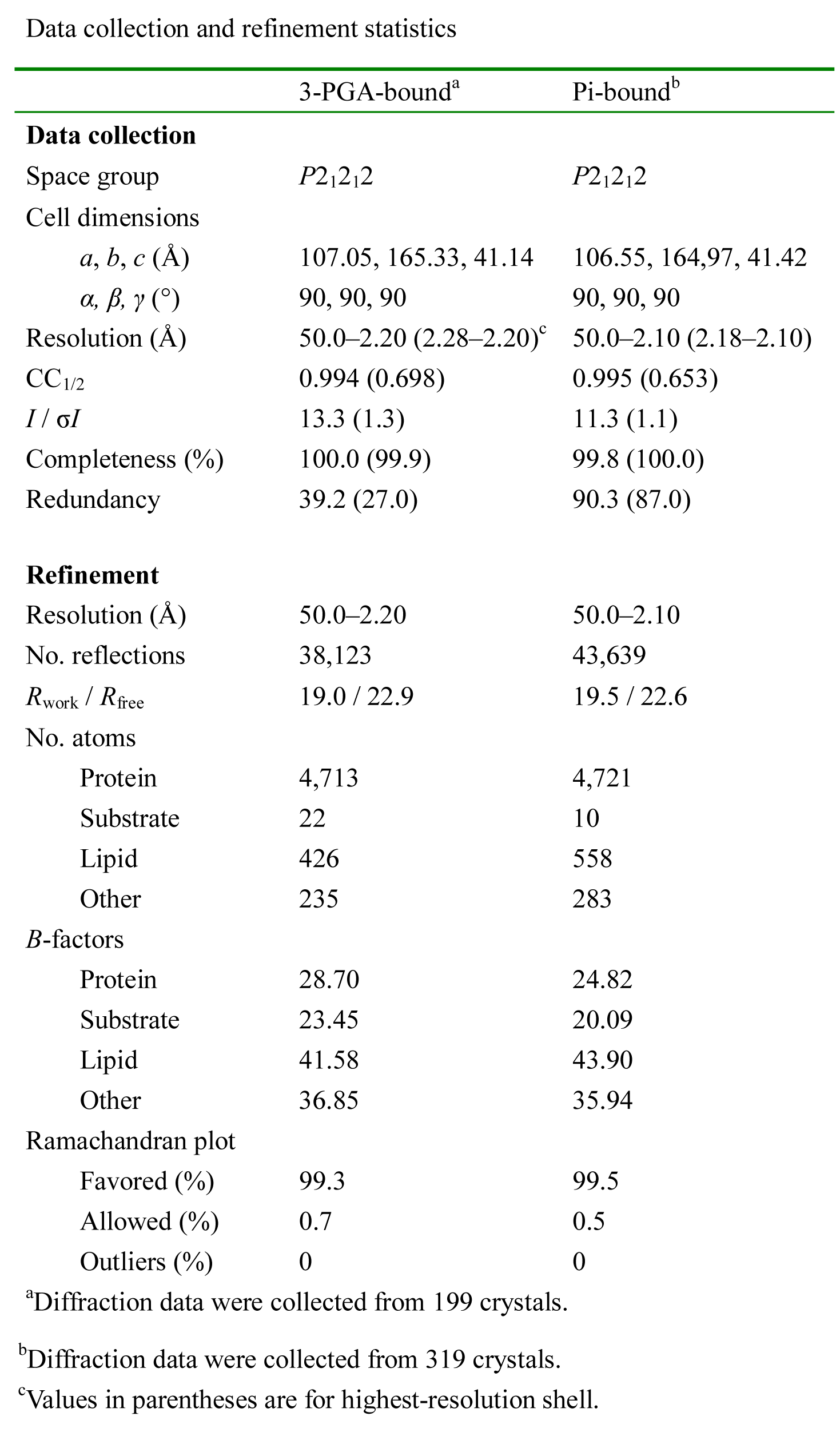
Data collection and refinement statistics

## Supplementary Videos

**Supplementary Video 1 | Conformational change of GsTPT2**

**Supplementary Video 2 | Molecular dynamics simulation of GsTPT2 without Pi**

